# Integrative proteomics and phosphoproteomics reveals phosphorylation networks involved in the maintenance and expression of embryogenic competence in sugarcane callus

**DOI:** 10.1101/2021.07.22.453415

**Authors:** Lucas R. Xavier, Felipe A. Almeida, Vitor B. Pinto, Lucas Z Passamani, Claudete Santa-Catarina, Gonçalo A. de S. Filho, Brian P. Mooney, Jay J. Thelen, Vanildo Silveira

## Abstract

Sugarcane (*Saccharum* spp.) is one of the most important crops for sugar, biofuel, and bioenergy production and has become an important commodity in the worldwide agricultural market in more than 100 countries. In this study, label-free quantitative proteomics and phosphoproteomics analyses were performed to investigate signaling events related to somatic embryo maturation and differentiation in sugarcane. Embryogenic callus (EC) at multiplication (EC0) and after 14 days (EC14) of maturation were compared. The EC14/EC0 comparison found that 251 phosphoproteins and 700 proteins were differentially regulated and accumulated, respectively. Metabolic pathway analysis showed that these proteins and phosphoproteins were enriched in lysine degradation and starch/sucrose metabolism during multiplication, whereas the differentiation of somatic embryos was found to involve the regulation of energetic metabolism, including the TCA cycle, oxidative phosphorylation, and carbon metabolism. Multiplication-related phosphoproteins were mainly associated with abscisic acid responses and transcriptional regulation of the TOPLESS (TPL), SNF1 kinase homolog 10 (KIN10), SEUSS (SEU), and LEUNIG_HOMOLOG (LUH) proteins. Among the maturation-related phosphoproteins, the phosphorylation of light harvesting complex photosystem ii, CURVATURE THYLAKOID 1B, vacuolar proton ATPase A1 and phytochrome interacting factor 3-LIKE 5 was found to be associated with bioenergetic metabolism and carbon fixation. A motif analysis revealed 15 phosphorylation motifs, and among these, the [D-pS/T-x-D] motif was unique among the phosphopeptides identified during somatic embryo differentiation. A coexpression network analysis of proteins and phosphoproteins revealed interactions among SNF1-related protein kinase 2 (SnRK2), abscisic acid responsive elements-binding factor 2 (ABF2), and KIN10, which indicated the role of these proteins in embryogenic competence in EC0. The interactions among ubiquitin-conjugating enzyme 5, ubiquitin-conjugating enzyme 35, small ubiquitin-like modifier 1, and histone deacetylase 1 may be involved in posttranslational protein modification during embryo maturation. Argonaute 1 (AGO1) also interacts with POLTERGEIST (POL) and may integrate gene silencing with the regulation of meristem identity during somatic embryo development. These results reveal novel dynamics of protein regulation in somatic embryogenesis and identify new potential players in somatic embryo differentiation and their phosphosites.

## 1. Introduction

Sugarcane (*Saccharum* spp.) is one of the most important crops for sugar, biofuel and bioenergy production and has become an important commodity in the worldwide agricultural market in more than 100 countries in tropical and subtropical regions (Waclawovsky et al., 2010). The genetic improvement of this crop is impaired by the polyploidy and aneuploidy (100-130 chromosomes) nature of the hybrids, which are mostly grown by conventional vegetative propagation due to the infertility and low viability of the seeds (Arruda, 2012; Garsmeur et al., 2018). Genetic transformation using *in vitro* embryogenic cells is one of the biotechnological alternatives for the development of cultivars with resistance traits and productivity but relies on the establishment of tissue culture protocols (Arruda, 2012). In this context, several molecular aspects involved in the regulation of *in vitro* plant development, including the mechanisms underlying the maintenance of embryogenic cell identity and the differentiation of these cells, remain unknown (Pierre-Jerome et al., 2018). The expression of embryogenic identity genes induces the ectopic formation of embryos and emblings, which represents a strategy for the regeneration of recalcitrant genotypes without major changes in the *in vitro* culture conditions.

Somatic embryogenesis is an *in vitro* culture approach that comprises the use of exogenous and/or endogenous signals to initiate autonomous embryo development in plant somatic cells (Fehér, 2015). Somatic embryogenesis efforts have been applied in the mass propagation and regeneration of transgenic plants of species of agronomic interest, including sugarcane (Lowe et al., 2016). The genetic pathways that lead to the differentiation of plant somatic cells into the embryogenic state and the expression of its embryogenic potential have been the focus of research on the molecular control of somatic embryogenesis (Fehér, 2015).

During the differentiation of plant cells, changes in gene expression modify the protein composition of the cells, resulting in an alteration in competence in response to signals (Pierre-Jerome et al., 2018). Gel-free proteomic studies have been performed to address key proteins during the development and differentiation of somatic embryos (Heringer et al., 2018). In sugarcane, 14-3-3, arabinogalactan, germin-like, heat-shock, IAA-amido synthase, late embryogenesis abundant, peroxidase, catalase, 6-phosphogluconate dehydrogenase, histone and ribosomal proteins are regulated during somatic embryogenesis (Heringer et al., 2015; Heringer et al., 2018; Reis et al., 2016). In addition, the proteins P-H^+^-ATPase and H^+^-PPase were quantified in microsomal fractions of sugarcane embryogenic callus (EC) and were found to be related to embryogenic competence (Passamani et al., 2018). The identification of these markers reveals new targets for functional genetics studies and will provide an important source of candidate genes to increase embryogenic competence.

Phosphorylation has been studied during somatic embryogenesis and represents a key step toward revealing the control and regulation of gene expression in this process (Almeida et al., 2020; Heringer et al., 2018). The phosphorylation of embryonic accumulated protein 1 (OsEM) and a B3 DNA binding domain-containing protein viviparous homolog (OsVP1) in callus tissue may be involved in hormonal signaling embryogenesis in rice (Wang et al., 2017). In *Elaeis guineensis* (oil palm), the use of phosphoproteomics at different stages of somatic embryogenesis allowed the identification of 460 phosphoproteins during the induction of embryogenic competence and the development of somatic embryos, and these phosphoproteins include an E3 ubiquitin-protein ligase (Aroonluk et al., 2020). Recently, 1279 phosphoproteins were identified by comparing ECs and nonembryogenic callus (NEC) in sugarcane (Almeida et al., 2020). It was possible to associate embryogenic competence with the phosphorylation of regulatory proteins such as ABA-responsive element binding factor 1 (ABF1), histone deacetylases 6 and 9 (HDA6 and 9), TOPLESS (TPL), argonaute 1b (AGO1) and SNF1 kinase homolog 10 (KIN10), a subunit of SnRK1 (Almeida et al., 2020). Together with the proteins regulated during somatic embryogenesis, phosphoproteins serve as potential candidate targets in translational proteomics research and agronomic purposes.

Here, we present a high-throughput label-free comparative phosphoproteomics and proteomics analysis of sugarcane embryogenic callus (EC) during the multiplication and maturation steps to identify new proteins and phosphoproteins associated with somatic embryo differentiation and morphogenesis. These results reveal novel dynamics of protein regulation in somatic embryogenesis and identify new potential players in somatic embryo differentiation and their phosphosites.

## 2. Materials and methods

### 2.1. Callus induction and somatic embryo maturation

EC induction and somatic embryo maturation were performed as previously described (Passamani et al., 2018). Briefly, for somatic embryo maturation three ECs (300 mg of fresh mass -FM-each) were placed in Petri dishes (90 × 15 mm) containing 20 mL of MS culture medium supplemented with 30 g L^-1^ sucrose, 2 g L^-1^ Phytagel and 1.5 g L^-1^ activated charcoal (Sigma-Aldrich). After inoculation, the cultures were maintained in a growth chamber at 25 ± 1 °C and grown in the dark for the first 7 days. Thereafter, the cultures were grown at a temperature of 25 ± 1 °C, and a 16-h photoperiod was established with GreenPower TLED 20-W W_m_B (Koninklijke Philips Electronics NV, Amsterdam, Netherlands) at 55 μmol m^-2^ s^-1^ for up to 42 days of culture. Samples of ECs at the multiplication phase and just before replacement with the maturation treatment (named EC0) and ECs after 14 days of incubation in the maturation treatment (named EC14) were collected for proteomics and phosphoproteomics analyses.

### 2.2. Protein extraction and digestion

Protein extraction and digestion for proteomics and phosphoproteomics analyses were performed according to Almeida et al. (2020). Five biological replicates (300 mg of fresh matter per sample) of EC0 and EC14 samples were lyophilized and ground using micropestles inside 1.5-mL microtubes. Subsequently, 1 mL of extraction buffer consisting of 7 M urea, 2 M thiourea, 2% (v/v) Triton X-100, 1% (w/v) dithiothreitol (DTT; GE Healthcare, Piscataway, USA), and 10 μL of EDTA-free Halt protease and phosphatase inhibitor cocktail (Thermo Fisher Scientific, Waltham, USA) was added to the sample powder. The protein concentration of each of the five biological replicates was estimated using a Bradford protein assay kit (Bio-Rad Laboratories, Hercules, USA).

Tryptic digestion was performed using the filter-aided sample preparation (FASP) method as described previously (Wiśniewski et al., 2009) with 1600 μg of protein from each biological replicate of the EC0 and EC14 treatments. Briefly, each biological replicate was divided into four aliquots of 400 μg due to the loading limit of the filter unit for tryptic digestion, and FASP was performed according to Almeida et al. (2020). The peptides from the four aliquots of each biological replicate were pooled, vacuum-dried, and resuspended in 800 μL of a solution containing 5% (v/v) acetonitrile (Thermo Fisher Scientific) and 0.1% (v/v) formic acid in mass spectrometry (MS)-grade water (Sigma-Aldrich). The peptide concentration was estimated by measuring A_205 nm_ using a NanoDrop 2000c spectrophotometer (Thermo Fisher Scientific). The peptides were stored at -80 °C prior to analyses.

### 2.3. Phosphopeptide enrichment

A High-Select™ TiO_2_ phosphopeptide enrichment kit (Thermo Fisher Scientific) was used for the enrichment of phosphopeptides from tryptic peptides. Aliquots of 500 µg of digested peptides from each biological replicate were enriched according to the manufacturer’s instructions. After elution of the phosphopeptides, the eluate was vacuum-dried and suspended in 25 μL of a solution containing 5% (v/v) acetonitrile and 0.1% (v/v) formic acid in MS-grade water. The phosphopeptide concentration was estimated by measuring A_205 nm_ using a NanoDrop 2000c spectrophotometer (Thermo Fisher Scientific). The phosphopeptides were stored at -80 °C prior to analyses.

### 2.4. Proteomics analysis

Proteomics analysis was performed using a nanoACQUITY ultraperformance liquid chromatograph (UPLC) coupled to a Q-TOF SYNAPT G2-Si instrument (Waters, Manchester, UK) with the parameters described by Botini et al. (2021). Briefly, the runs consisted of five biological replicates; in each run, 1 μg of the digested proteins were loaded onto a nanoACQUITY UPLC M-Class Symmetry C18 5 μm trap column (180 μm × 20 mm) at 5 μL min^-1^ for 3 min and then onto a nanoACQUITY M-Class HSS T3 1.8 μm analytical reversed-phase column (75 μm × 150 mm) at 400 nL min^-1^. The column temperature was set to 45 °C. For peptide elution, a binary gradient was used, and mobile phases A and B consisted of water (Tedia, Fairfield, USA) and 0.1% formic acid and acetonitrile (Sigma-Aldrich) and 0.1% formic acid, respectively. Mass spectrometry was performed in the positive and resolution mode (V mode) with ion mobility (HDMS^E^) and in the data-independent acquisition (DIA) mode.

### 2.5. Phosphoproteomics analysis

Phosphoproteomics analysis was performed using a nanoElute (Bruker Daltonics, Bremen, Germany) nanoflow chromatography system coupled to a Q-TOF timsTOF Pro (Bruker Daltonics) according to Almeida et al. (2020). Briefly, the runs consisted of five biological replicates of 1 μg of enriched phosphopeptides loaded onto a C18 trap column (Thermo Fisher Scientific; PepMap 100, 5 cm x 300 μm) and then separated using a 70-min gradient at 400 μL min^-1^ with a pulled-needle, self-packed, fused silica analytical column (containing Waters BEH-C18, 1.7-μm particles, 20-cm length x 75-μm inner diameter). The column temperature was 50 °C. For phosphopeptide elution, a binary gradient was used: mobile phase A consisted of MS-grade water and 0.1% formic acid, and mobile phase B consisted of 100% acetonitrile and 0.1% formic acid. MS and MS/MS data were acquired using the parallel accumulation–serial fragmentation (PASEF) method (Meier et al., 2015; Meier et al., 2018).

### 2.6. Proteomics and phosphoproteomics data analysis

Both proteomics and phosphoproteomics MS/MS spectra were searched against the protein databank generated from the sequencing of the haploid *Saccharum spontaneum* AP85-441 (databank version: v20190103). The proteomics and phosphoproteomics MS data have been deposited in the ProteomeXchange Consortium via the PRIDE (Perez-Riverol et al., 2019) partner repository with the dataset identifier PXD027388.

The proteomics MS/MS spectra processing and database search were performed using ProteinLynx Global SERVER (PLGS) software v.3.02 (Waters), and label-free quantification analyses were performed using ISOQuant software v.1.7 (Distler et al., 2014) with previously described parameters (Passamani et al., 2019). To ensure the quality of the results after data processing, only proteins present in at least four runs or absent in all biological replicates (for unique proteins) were considered for differential accumulation analysis in the EC14/EC0 comparison. Proteins with significant Student’s t-test (two-tailed; P < 0.05) results were considered differentially accumulated (DAP), considering up-accumulated if the Log_2_ fold change (FC) was greater than 0.5 and down-accumulated if the log_2_ FC was less than -0.5.

The phosphoproteomics MS/MS spectra processing and database search were performed using MaxQuant software v. 1.6.3.4 (Cox and Mann, 2008) as described in (Almeida et al., 2020). After MaxQuant data analyses, the resulting table of identified phosphosites (phospho STY (sites).txt) was filtered only for those that were confidently localized (localization probability ≥ 0.75). Only the proteins containing high-confidence phospho (STY)-modified peptides and that exhibited LFQ intensity in at least four biological replicates or were absent in all biological replicates (for unique proteins) were considered for differential regulation analysis in the EC14/EC0 comparison. Phosphoproteins were considered differentially regulated (DRP) based on their Student’s t-test results (two-tailed; P < 0.05); a phosphoprotein was considered up-regulated if its Log_2_ FC value was greater than 0.5 and down-regulated if the Log_2_ FC value was less than -0.5.

Functional annotation analysis was performed using OmicsBox software (https://www.biobam.com/omicsbox) and UniProtKB (https://www.uniprot.org). The protein sequences were submitted to a BLAST search against the NCBI nonredundant green plant protein database (taxa: 33090, Viridiplantae). An analysis of the enrichment of biological processes, cellular components and molecular functions of DAPs and DRPs was performed by Fisher’s exact test (P < 0.01) using OmicsBox. A Kyoto Encyclopedia of Genes and Genomes (KEGG) enrichment pathway analysis between DAPs and DRPs (P < 0.01) was performed in Metascape (Zhou et al., 2019) after a BLAST search of NCBI (https://www.ncbi.nlm.nih.gov) to obtain *Arabidopsis thaliana* reference sequences.

The interaction networks of DAPs and DRPs were constructed using *Arabidopsis* homologs of sugarcane identified through a STRING search followed by the downstream analysis in Cytoscape (version 3.7.1) (Shannon et al., 2003). The predicted kinases, transcription factors and transcriptional regulators among DAPs and DRPs were identified using iTAK software (Zheng et al., 2016).

MotifeR software was used to identify possible phosphorylation motifs related to the differentiation of sugarcane embryogenic cells (Wang et al., 2019). The identification of motifs was performed considering a minimum of 20 phosphopeptides and a P-value threshold of < 0.000001. The sugarcane proteome and phosphoproteome established in this study served as the background, the width was set to 13, and the number of occurrences was set to 20.

## 3. Results

### 3.1. Somatic embryo differentiation and development

ECs after 14 days of maturation treatment (EC14) showed the onset of somatic embryo differentiation (Supplementary Fig. S1B). After this period, differentiation and development occurred simultaneously, resulting in the formation of mature somatic embryos after 42 days of maturation treatment (Supplementary Fig. S1C). No signs of somatic embryo maturation were observed in the ECs during multiplication (EC0) (Supplementary Fig. S1A).

### 3.2. Comparative label-free proteomics and phosphoproteomics analysis of EC during multiplication and maturation of somatic embryos

The proteomics analysis identified 1620 proteins in the ECs at maturation compared with those from the multiplication treatment (EC14/EC0 comparison), and a total of 699 DAPs were obtained (Fig. 1A; Supplementary Table S1). Among the DAPs, 339 and 297 proteins were up- and down-accumulated, respectively, and 29 and 34 proteins were uniquely found in EC0 and EC14, respectively.

**Fig. 1.**
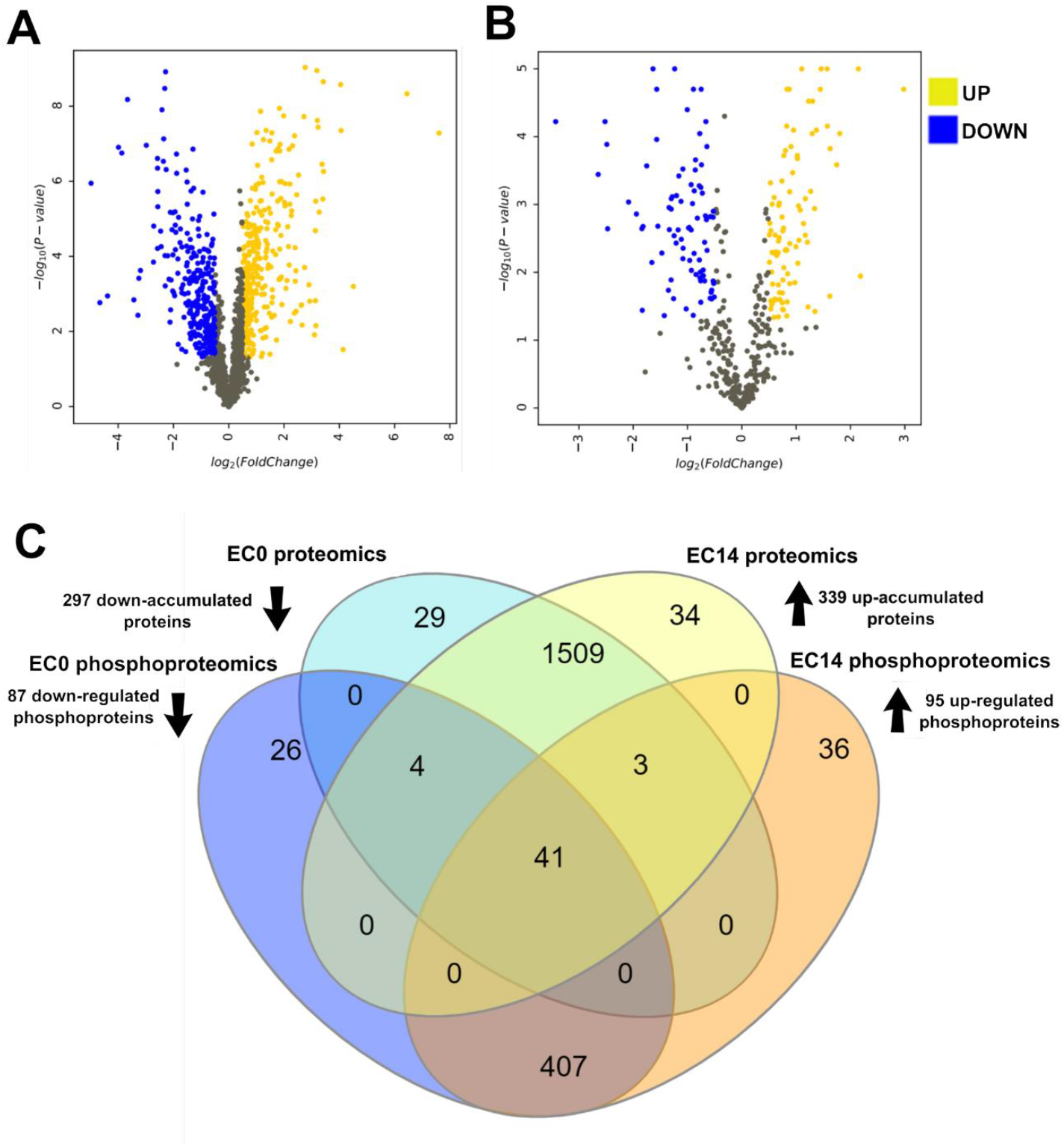
Comparative proteomics and phosphoproteomics of ECs at maturation compared with EC at the multiplication phase before maturation (EC14/EC0 comparison). Volcano plot of differentially accumulated proteins (DAPs) (A) and differentially regulated phosphoproteins (DRPs) (B) identified from the EC14/EC0 comparison. The yellow and blue points represent significant observations (q value < 0.05). Venn diagram of protein and phosphoprotein IDs identified in EC0 and EC14 (C).

The phosphoproteomics analysis allowed the identification of 1471 phosphoproteins, including 251 DRPs, from the EC14/EC0 comparison (Fig. 1B; Supplementary Table S2). Of the DRPs, 95 and 87 were up- and down-regulated, respectively, and 39 and 30 unique phosphoproteins were identified in EC14 and EC0, respectively (Fig. 1B).

To address the impact of protein phosphorylation on somatic embryo differentiation, a high-throughput phosphoproteomics analysis was performed based on the EC14/EC0 comparison. A total of 1704 phosphopeptides were identified, and these resulted in the discovery of 1845 phosphosites (phosphorylation probability > 0.75) (Supplementary Table S3). Most of the identified phosphosites corresponded to serine (S) residues (1689 sites, 92%), followed by threonine (T) residues (146 sites, 8%) and tyrosine (Y) residues (10 sites, < 1%) (Supplementary Fig. S2A). An analysis of the number of phosphosites on each phosphopeptide revealed that the majority had only one phosphosite (1566), followed by two (132), three (1) and four (1) phosphosites (Supplementary Fig. S2B).

To reveal the pathways of protein regulation during sugarcane somatic embryo morphogenesis, label-free proteomics and phosphoproteomics analyses were performed by comparing ECs at 14 days of maturation (EC14) with ECs from the multiplication phase and just before the maturation treatment (EC0) (EC14/EC0 comparison) (Supplementary Fig. S1).

Regarding the overlap in the proteomics and phosphoproteomics data, 7.14% of the sugarcane protein IDs were identified simultaneously in the proteomics and phosphoproteomics analysis (Fig. 1C). A total of 30 DRPs were also found as DAPs (Supplementary Fig. S3). After functional evaluation and a search of the databases and literature, 19 DRP phosphoproteins with a putative role in embryogenic competence and the differentiation of somatic sugarcane embryos were identified (Table 1).

**Table 1.**
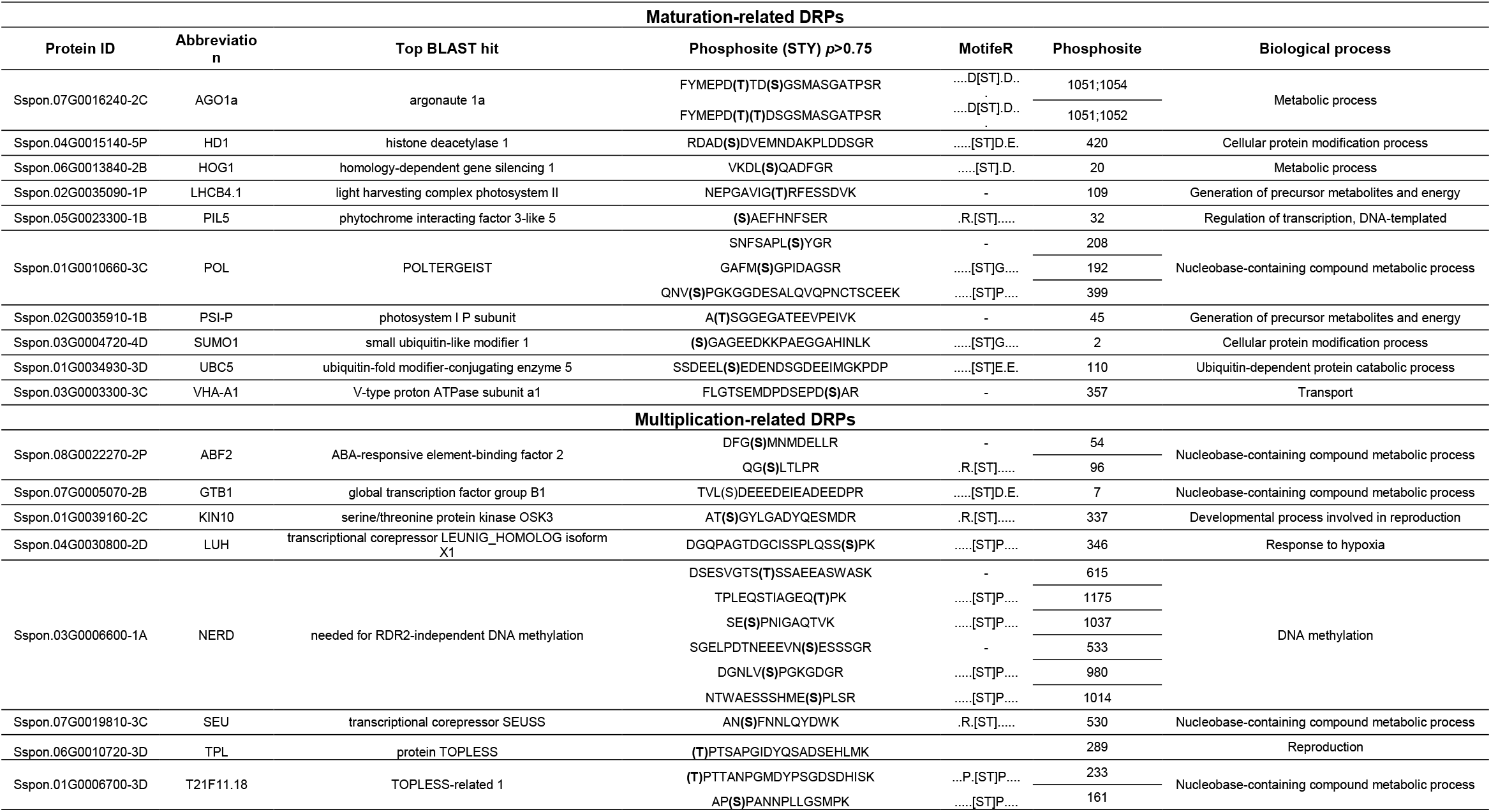
Phosphoproteins with putative roles in embryogenic competence and their abundance during somatic embryogenesis of sugarcane identified by comparing ECs from the maturation treatment (EC14) to those at multiplication (ECO) (EC14/EC0 comparison).

### 3.3. Functional annotation and enrichment of metabolic pathways related to DAPs and DRPs

The analysis of the maturation-related biological processes identified a large number of DAPs and DRPs involved in photosynthesis, response to cytokinin, translation, oxidation-reduction process and generation of precursors for metabolites and energy (Fig. 2A). In contrast, the multiplication-related DAPs and DRPs were mainly grouped in protein metabolic processes, responses to stress catabolic processes and responses to abiotic stimuli (Fig. 2A). A gene ontology enrichment showed that cytoplasm, chloroplast and plastid localization and molecular functions of catalytic activity and enzyme regulator activity were enriched in maturation-related DAPs and DRPs (Supplementary Figs. S4-S5). The multiplication-related DAPs and DRPs were found to be related to the regulation of nucleic acid-templated transcription, transcription regulator activity, DNA-templated transcription, RNA biosynthetic process, DNA binding transcription factor activity, and positive regulation of RNA metabolic process (Supplementary Figs. S4-S5).

**Fig. 2.**
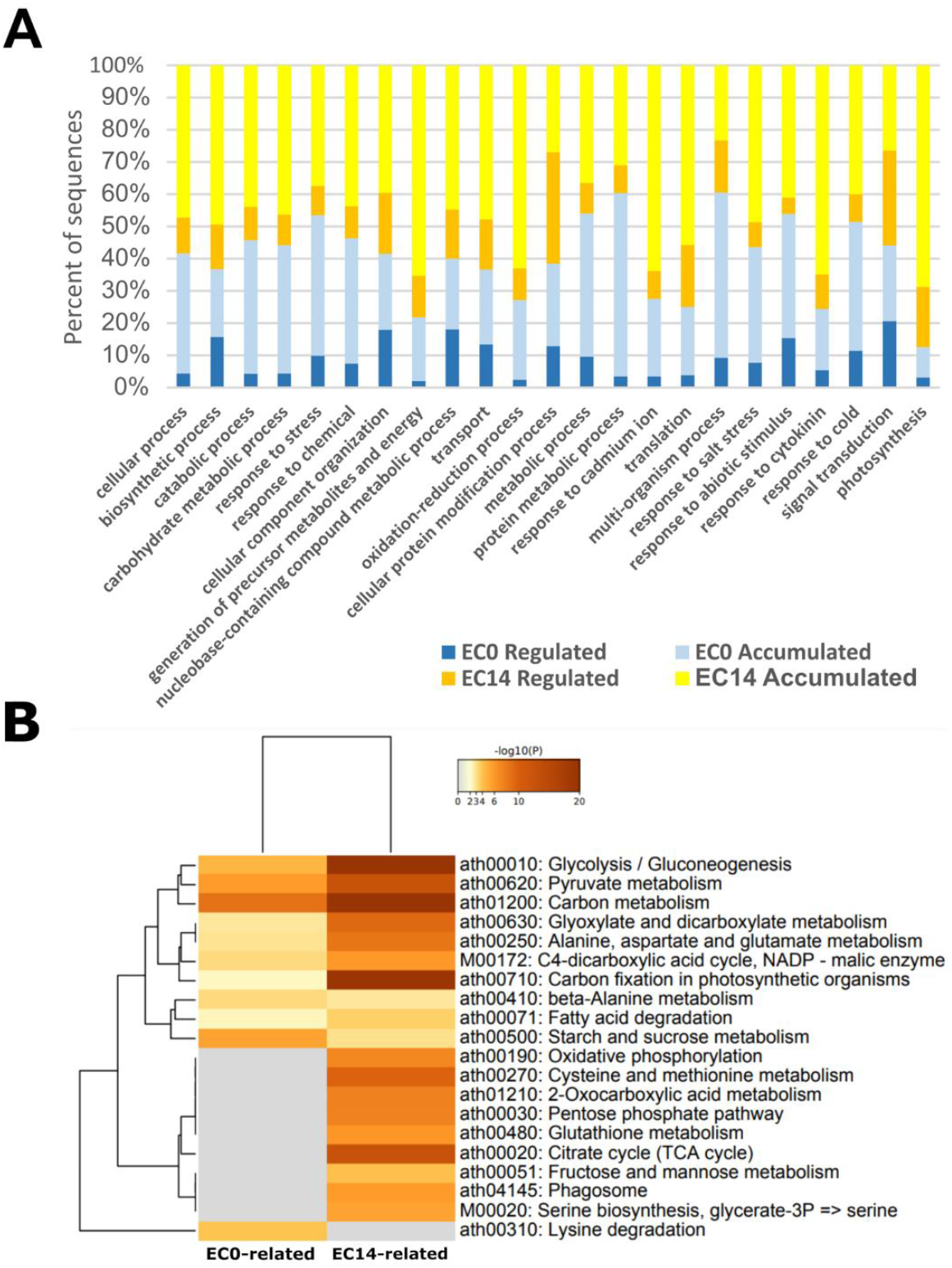
Functional annotation and enrichment of metabolic pathways related to DAPs and DRPs. (A) Evaluation of the most representative biological processes among DAPs and DRPs. (B) Heatmap of KEGG pathway enrichment analysis (p < 0.01) of the multiplication- and maturation-related DAPs and DRPs.

The KEGG pathway enrichment analysis clustered the DAPs and DRPs of each group together among maturation-related and multiplication-related proteins (Fig. 2B). The maturation-related proteins were enriched in glycolysis/gluconeogenesis, carbon metabolism, and carbon fixation in photosynthetic organism pathways. Fructose and mannose metabolism, the TCA cycle, glutathione metabolism, the pentose phosphate pathway and oxidative phosphorylation were enriched only in the maturation-related DAPs and DRPs. In the multiplication-related groups, starch and sucrose metabolism were enriched, whereas the lysine degradation pathway was unique to multiplication-related DAPs and DRPs (Fig. 2B).

### 3.4. Identification of enriched phosphorylation motifs and analysis of phosphosite conservation

A MotifeR analysis was performed to determine the overrepresented phosphorylation motifs in the phosphopeptides. A total of 15 motifs were found to be distributed among 76.2% of the sequenced phosphopeptides and occurred in both multiplication- and maturation-related DRPs (Fig. 3A). The [pS/T-D-x-E] motif (DS subtype enriched in the nucleus), which comprises transcription-related proteins such as the transcription elongation factor SPT6-like X1 (GTB1), DEAD-box ATP-dependent RNA helicase 42, putative histone deacetylase 19 (HD1), and CCR4-NOT transcription complex subunit 10, and the motif [pS/T-P-x-R] (basic SP-like motif), which includes the phosphoproteins DA1-related 1-like protein, TOM1-like protein 1, and ubiquitin carboxyl-terminal hydrolase 36, were the most representative of maturation-related DRPs (Fig. 3A). In contrast, the [pS/T-F] motif (target of OXI1) (Fig. 3A) was the most representative of multiplication-related DRP, including GEM-like protein 1 and cell division cycle 5-like protein, which regulates development and the cell cycle, respectively. No motifs with phosphorylated threonine (T) or tyrosine (Y) were found.

**Fig. 3.**
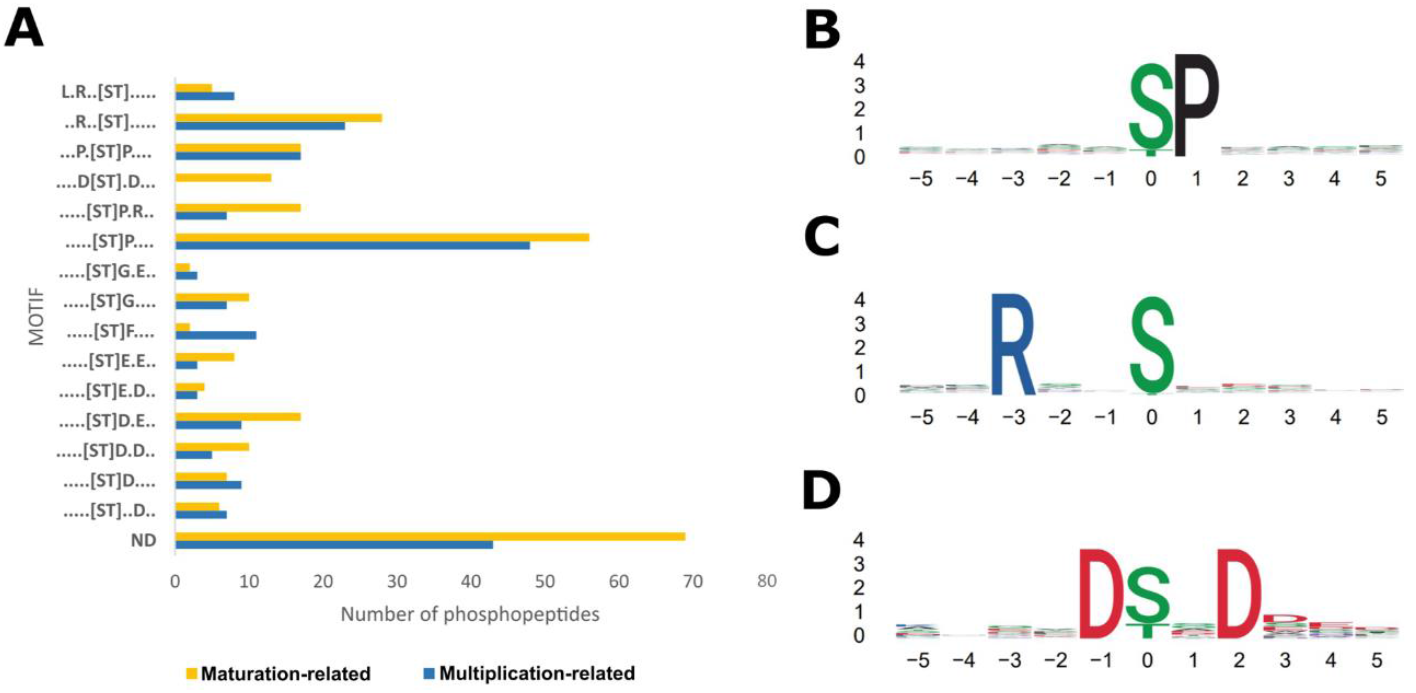
Identification of enriched phosphorylation motifs. (A). All phosphorylation motifs identified after enrichment analysis using MotifeR and the comparison between DRPs. (B-D). Motifs overrepresented in the phosphopeptides.

The [pS/T-P] motif (Fig. 3B) was the most representative among all the identified phosphopeptides, including regulatory phosphoproteins, such as the transcriptional corepressor LEUNIG_HOMOLOG X1 (LUH). The [R-x-x-pS/T] motif was the second most representative (Fig. 3C) and was found in phosphoproteins related to the ethylene response, such as AP2-like ethylene-responsive transcription factor AIL1 (AIL1) and auxin-responsive protein SAUR76-like, in addition to those with catalytic activity, such as glucose-6-phosphate 1-dehydrogenase (G6PD6), soluble inorganic pyrophosphatase (PPa3), and phosphoenolpyruvate carboxylase 2 (PPC2). The [D-pS/T-x-D] motif (core-enriched DS subtype) was present only in the phosphopeptides identified in maturation-related DRPs (Fig. 3D), including the phosphoprotein argonaute 1A (AGO1a) and 26S proteasome regulatory subunit RPN13, which regulate gene silencing and protein degradation, respectively.

The phosphosite conservation analysis showed 96% and 65% conservation of phosphosites in *Zea mays* and *Arabidopsis*, respectively. The average identity between the sugarcane phosphoproteins and maize sequences was 86%, which is higher than the average of 62% found between sugarcane and *Arabidopsis* phosphoproteins (Supplementary Table S4).

### 3.5. Kinase activity and transcription factor and regulator identification

The annotation of proteins with kinase activity among DAPs and DRPs resulted in the identification of 28 proteins (Supplementary Table S5). Among the 13 kinases related to multiplication, three serine/threonine protein kinases, OSK3, cell division cycle 5-like protein and CCR4, were highlighted. Among the maturation-related proteins, 15 protein kinases were found, and these included the nuclear pore complex protein NUP1 and the putative serine/threonine protein kinase IREH1. In addition, 22 transcription factors were found among DAPs and DRPs, and these included 18 multiplication- and four maturation-related factors (Supplementary Table S5). The four maturation-related transcription factors were the transcription factor BIM3, the protein SPATULA, the single-stranded DNA-binding protein WHY2 and zinc finger CCCH domain-containing protein 40. The regulated multiplication-related transcription factors included the AP2-like ethylene-responsive transcription factor AIL1, dr1-associated corepressor, probable WRKY transcription factor 32 and abscisic acid responsive element-binding factor 2 (Supplementary Table S5). In addition, four multiplication-regulated transcriptional regulators, including LUH and H3 lysine-9 specific SUVH1 (SUVH1), were also found among both the DAPs and DRPs (Supplementary Table S5). The maturation-related transcriptional regulators enhancer of zeste 3 and DA1-related 1-like protein were identified (Supplementary Table S6).

### 3.6. Protein-protein interaction networks between proteins and phosphoproteins

To visualize the interactions between the accumulated and regulated phosphoproteins, a coexpression network with confidence > 0.7 was constructed with potential key phosphoproteins involved in the expression of embryogenic competence (Fig. 4, Table 1). Using the *Arabidopsis* databank, the interaction among SNF1-related protein kinase 2.6 (SnRK2.6), abscisic acid responsive elements-binding factor 2 (ABF2), and SNF1 kinase homolog 10 (KIN10) was observed to be regulated as multiplication-related DRPs. The interaction among the proteins TOPLESS (TPL), histone deacetylase 1 and TOPLESS-related 1 (TPR1, shown in Fig. 4 as T21F11.18) was also detected (Fig. 4).

**Fig. 4.**
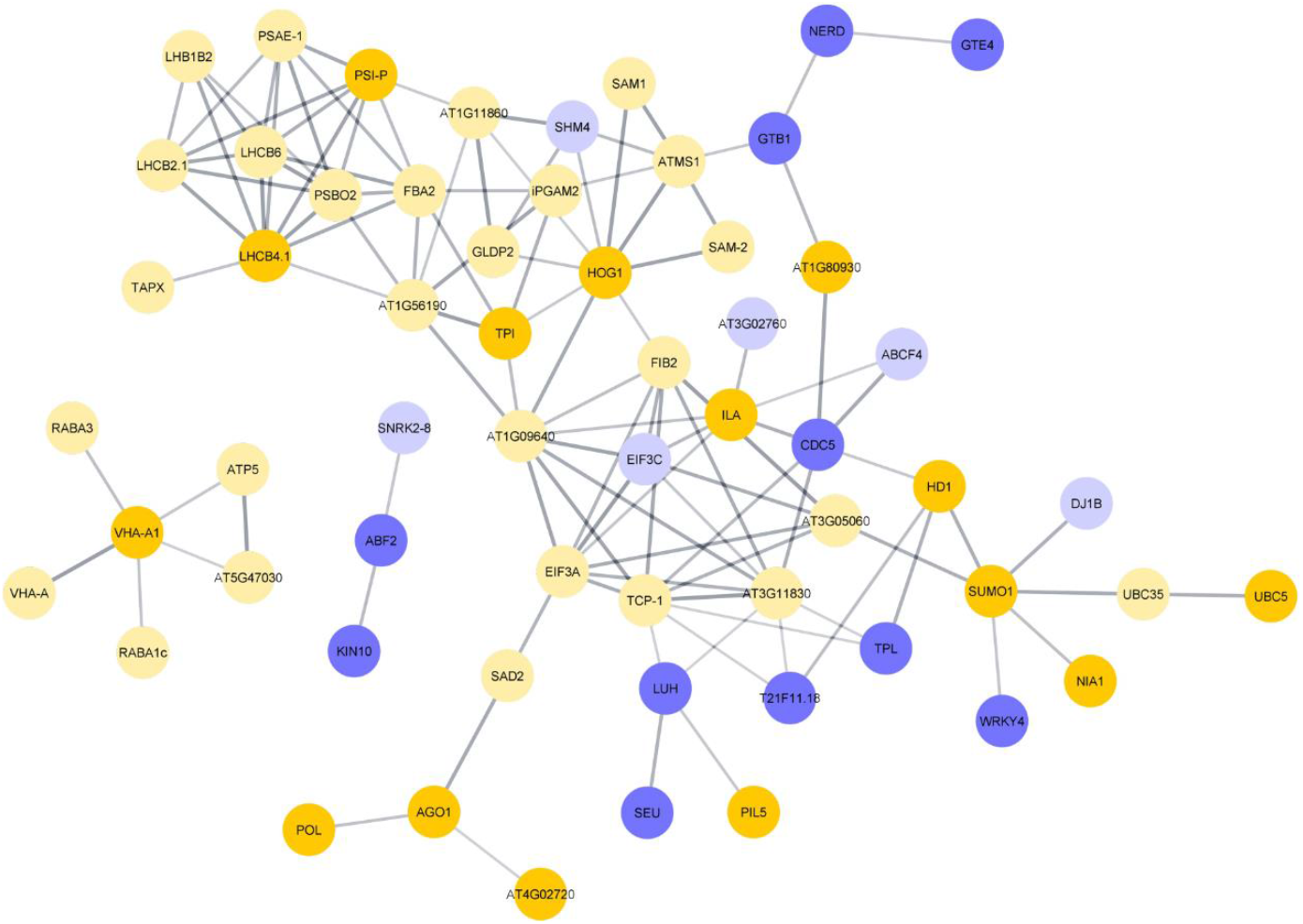
Protein-protein interaction network between DAPs and DRPs with putative roles in embryogenic competence. The analysis was performed using *Arabidopsis* orthologs identified by STRING using the sugarcane amino acid sequence. Only first neighbors are shown in the interactions. The dark and light blue ellipses represent multiplication-related DAPs and DRPs, respectively, and the dark and light-yellow ellipses represent maturation-related DAPs and DRPs, respectively.

Interactions among maturation-related DAPS proteins involved in posttranslational modification, such as ubiquitin-conjugating enzyme 5 (UBC5), ubiquitin-conjugating enzyme 35 (UBC35), small ubiquitin-like modifier 1 (SUMO1), and histone deacetylase 1 (HD1), were observed during embryo maturation. AGO1 also interacts with POL during somatic embryo development. In addition, proton pump vacuolar proton ATPase A1 (VHA-A1) interacts with the delta subunit of mt ATP synthase (ATP5), RAB GTPase homolog A1C (RABA1c), RAB GTPase homolog A3 (RABA3), vacuolar ATP synthase subunit A (VHA-A) and AT5G47030, a mitochondrial ATP synthase subunit delta (Fig. 4).

The interactions among light harvesting complex photosystem II (LHCB4.1), curvature thylakoid 1B (PSI-P), fructose-bisphosphate aldolase 2 (FBA2), photosystem ii subunit O-2 (PSBO2), subunit E of photosystem I (PSAE-1), light-harvesting complex photosystem II subunit 6 (LHCB6), light-harvesting chlorophyll B-binding 2 (LHCB2. 1) and thylakoidal ascorbate peroxidase (TAPX), members of the photosynthetic apparatus, were detected as maturation-related DRPs (Fig. 4).

During the development of somatic embryos, multiplication-related homology-dependent gene silencing 1 (HOG1) was found to interact with S-adenosylmethionine synthetase 1 and 2 (SAM1 and SAM2), cytosolic isoform triose phosphate isomerase (TPI) and serine hydroxymethyltransferase 4 (SHM4) (Fig. 4). The cell division cycle 5 (CDC5) protein interacts with ATP-binding cassette F4 (ABCF4) as a multiplication-related phosphoprotein and with histone deacetylase 1 (HD1) and ilithyia (ILA) as maturation-related proteins (Fig. 4). Among multiplication-related proteins, global transcription factor group b1 (GTB1), global transcription factor group e4 (GTE4) and rdr2-independent DNA methylation (NERD) also interact in the network (Fig. 4).

Due to the central regulatory role of calcium-dependent protein kinases (CPKs) and cyclin-dependent kinases (CDKs) in signal transduction, an interaction network between CPKs and CDKs identified among DAPs and DRPs was also constructed (Supplementary Fig. S6; Supplementary Table S4). Interactions among calcium-dependent protein kinases 4, 5, 6 and 20 (CPK4, CPK5, CPK6 and CPK20), ascorbate peroxidase 3 (APX3), plasma membrane proton ATPase (HA1), and root hair defective 2 (RHD) (Supplementary Fig. S6) were observed. Among the CDKs, only CDK5 was found to interact with other proteins (Supplementary Fig. S6). CDK5 interacts with 8 down-regulated proteins and 5 down-regulated phosphoproteins, including LHP1-interacting factor 2 (LIF2), regulatory particle AAA-ATPase 2A (RPT1A) and heat shock protein 70 (HSP70). Among the maturation-related DAPs and DRPs that interact with CDC5, proteins involved in transcription and gene expression, such as ARGININE/SERINE-RICH 45 (SR45), eukaryotic translation initiation factor 4G (EIF4G), histone deacetylase 1 (HD1) and eukaryotic release factor 1-3 (ERF1-3), were identified. Maturation-related proteins involved with cytoskeletal tubulin alpha 2, 5 and 6 (TUA2, TUA5 and TUA6) were observed and interacted with CDC5 (Supplementary Fig. S6).

## 4. Discussion

### 4.1. The maintenance of embryogenic competence in multiplication is related to networks of repressor phosphoproteins

A challenge in plant developmental biology is understanding how the maintenance of stem cell identity is regulated and how these cells differentiate (Pierre-Jerome et al., 2018). As a result, we performed proteomics and phosphoproteomics analyses comparing sugarcane callus at the beginning (EC0) and after 14 days of maturation treatment (EC14) with the aim of obtaining a better understanding of the molecular basis underlying the regulation of somatic embryo maturation and differentiation.

Dormancy-associated gene-1/auxin-repressed protein (DRM1/ARP) was grouped into multiplication-related DRPs and phosphorylated at T58 and S84 (which correspond to T50 and S71 in *Arabidopsis*) (Supplementary Table 4). These phosphosites have been identified in other studies with *Arabidopsis* (Nukarinen et al., 2017; Reiland et al., 2009; Roitinger et al., 2015), which indicates their importance in the activity of DRM1. This protein is a marker for dormancy of plant meristems and belongs to the auxin-repressed superfamily (ARP) (Lee et al., 2013; Rae et al., 2013). In seeds and lateral buds, dormancy is characterized by reduced metabolic activities and insensitivity to signals that promote growth and development (Graeber et al., 2012). The presence of proteins involved in dormancy that are phosphorylated at EC0 (multiplication phase) indicates that the maintenance of embryogenic competence may involve negative regulatory processes at the posttranscriptional level.

WUS, a master regulator of somatic embryogenesis and plant meristem identity, reportedly acts as a transcriptional repressor to maintain the dedifferentiated state of apical meristem cells through TOPLESS (TPL) and TOPLESS-like repressor proteins (Jha et al., 2020). In our data, TPL and TOPLESS-related 1 (TPR1; T233) were phosphorylated in EC0 and interacted with HD1 (also known as HD19), which was phosphorylated at EC14 (Fig. 4). In maize, *RAMOSA1 ENCHANCER LOCUS 2* (REL) encodes a transcriptional corepressor orthologous to AtTPL and is able to rescue the embryonic defects of the *topless-1* mutant of *Arabidopsis* (Gallavotti et al., 2010). Furthermore, mutations in TPL-1 affect the apical-basal patterning of *Arabidopsis* embryos and result in the formation of two root poles (Long et al., 2006). This finding indicates that TPL activity must be regulated for the progression of embryogenesis. Because the induction of embryogenic potential involves a dedifferentiated cellular state, the maintenance of this potential can be linked to the repression of differentiation programs. The regulation of TPL and TPR1 in embryogenic competence may be a factor that indirectly inhibits the expression of the embryogenic program and promotes callus embryogenic competence under 2,4-D treatment.

In *Arabidopsis*, NERD performs transcriptional silencing of specific genes by integrating signals from chromatin and silencing mRNAs (Pontier et al., 2012). In EC0 cells, NERD interacts with GTE4 and GTB1 in the protein-protein network (Fig. 4). In GTE4 mutants, a loss of function of cells in the quiescent root center has been identified, which indicates that this gene participates in the activation and maintenance of cell division in meristems (Airoldi et al., 2009). In addition, germination is delayed in GTE4 mutants (Della Rovere et al., 2010). GTB1, also known as suppressor of Ty insertion 6-like (SPT6L), participates in apical-basal patterning in *Arabidopsis* embryos through negative regulation of PHABULOSA and PHAVOLUTA, likely through miRNA recruitment (Gu et al., 2012). Thus, the regulation of NERD, GTB1 and GTE4 in EC0 may be related to the transcriptional repression of genes involved in the somatic embryo differentiation program, which maintains the embryogenic cells in the multiplication phase.

In our analyses, LUH and SEU were classified as multiplication-related proteins that interact with PIL5, which was uniquely found at the maturation phase (EC14) (Fig. 4, Table 1). In the present study, the corepressor LUH was found to be phosphorylated at S346, which shares the same phosphosite as maize and corresponds to S345 in *Arabidopsis* (Supplementary Table 4). This phosphosite was identified during the early stages of maize development (Walley et al., 2016) and in ABA-treated *Arabidopsis* seedlings (Wang et al., 2013). LUH has been associated with the repression of homeotic genes linked to floral and embryogenic development (Sitaraman et al., 2008). LUH activity depends on the adaptor protein SEUSS (phosphorylated at S530), which form a corepressor complex that regulates floral cell specificity and identity by repressing the homeotic AGAMOUS gene (Franks et al., 2006). In *Arabidopsis*, the phosphorylation of PIL5, also known as phy-interacting factor 1 (PIF1), is signaled by light and leads to its degradation, which decreases PIL5 repression activity during photomorphogenesis (Xu et al., 2015). The interaction between LUH and PIF1 is already known and is directly involved in the inhibition of *Arabidopsis* seed germination (Lee et al., 2015). Thus, the interaction among PIL5, LUH and SEU is supposedly involved in the maintenance of the dedifferentiated state of ECs during multiplication (EC0). In this sense, the phosphorylation and light-mediated degradation of PIL5 in EC14 would signal that embryogenic cells progress to the differentiation of somatic embryos under maturation treatment.

### 4.2. Somatic embryo maturation progression involves the accumulation and phosphorylation of proteins related to bioenergetic metabolism

The production of chlorophyll is a visible aspect of the morphology of somatic embryos differentiating into callus at 14 days of maturation (EC14) (Supplementary Fig. S1). This observation indicates that the embryo photosynthetic apparatus is developing and that the embryos are becoming autotrophic. Accordingly, our proteomics and phosphoproteomics analysis revealed that maturation-related groups of proteins and phosphoproteins were mainly involved in carbon metabolism, pyruvate metabolism, glycolysis/gluconeogenesis, carbon fixation in photosynthetic organisms, and the TCA cycle (Fig. 2B). These data indicate that the accumulation and phosphorylation of proteins involved in photosynthesis is a feature of differentiating somatic embryos during maturation.

Several maturation-related DAPs and DRPs are involved in the carbon fixation process (Fig. 4; Table 1). Among them, LHCB4, phosphorylated at T109 in sugarcane and at corresponding T112 in *Arabidopsis* (Table 1), is a phosphosite that was identified in previous studies (Al-Momani et al., 2018; Nukarinen et al., 2017). Defective LHCB4 mutants of *Arabidopsis* reduce LHCII assembly and decrease the photosystem II quantum yield (Chen et al., 2018). LHCB4 is a monomeric antenna protein that binds light-harvesting complex (LHCII) trimers to photosystem II and is critical for the transfer of energy from sunlight to the reaction center during photosynthesis (van Bezouwen et al., 2017). PSI-P is phosphorylated at T45 in sugarcane (Table 1) and corresponds to T65 in *Arabidopsis*, which is phosphorylated during anther development (Ye et al., 2016). Phosphorylation of PSI-P depends on light intensity and is related to the integrity of photosystem II in *Arabidopsis*, where PSI-P proteins are also needed for growth (Pribil et al., 2018; Trotta et al., 2019). Thus, the regulation of and interaction among these proteins and phosphoproteins involved in photosynthesis may be related to the onset of photomorphogenesis observed in EC14 under maturation treatment (Supplementary Fig. S1).

The VHA-A1 phosphoprotein was up-regulated in the maturation-related DPRs (Table 1) and phosphorylated at S357, which corresponds to S689 in maize (Supplementary Table 4). This phosphorylated residue has previously been identified in pollen (Chao et al., 2016) and embryo and endosperm tissues of maize (Walley et al., 2016), which indicates a role for the phosphorylation of this residue during the initial stages of monocot development. In the protein-protein interaction network, VHA-A1 interacts with ATP5, RABA1c, RABA3, VHA-A and AT5G47030, a mitochondrial ATP synthase subunit delta (Fig. 4). This result is in accordance with the findings from the analysis of the maturation-related DAPs and DRPs involved in energy metabolism, which revealed enrichment of several carbon metabolism pathways (Fig. 2B). The interaction among these proteins and phosphoproteins may be related to the increased metabolic activity previously observed in sugarcane callus after 14 days of maturation (Passamani et al., 2018). Additionally, soluble inorganic pyrophosphatase (PPa3) was identified as unique and up-regulated in maturation-related DAPs and DRPs, respectively (Supplementary Tables S1-S2). PPa3 was phosphorylated at S38, which corresponds to S36 in maize and S28 in *Arabidopsis* (Supplementary Tab S4), and this protein was also identified in pollen tissues of maize (Chao et al., 2016) and *Arabidopsis* (Mayank et al., 2012). Thus, VHA-A1 and PPase activity has been demonstrated to be related to the metabolic pathways linked to the differentiation of sugarcane somatic embryos (Passamani et al., 2018), and our data suggest that the observed increase in proton pump activity may be related to their phosphorylation.

### 4.3. Insights into phosphoproteins involved in the epigenetic regulation of somatic embryogenesis

Epigenetic modifications play a central role in regulating gene expression through DNA methylation, chromatin remodeling, and small RNAs (Kumar and Van Staden, 2017). In somatic embryogenesis, the level of chromatin condensation leads to changes in gene expression and integrates hormonal, stress and developmental pathways during the activation of the embryogenic program in competent cells (Fehér, 2015). Both AGO1 and HD19 were found to be involved in epigenetic regulation and were differentially phosphorylated in EC14 compared with EC0 (Fig. 4). CDC5 regulates development and the cell cycle in *Arabidopsis* by accumulating miRNA and integrating the DICER-LIKE1 complex (Zhang et al., 2013). During somatic embryogenesis in sugarcane, CDC5 was found to interact with LIF2, RPT1A and HSP70 in EC0. In addition, CDC5 interacts with maturation-related DAPs and DRPs involved in transcriptional and epigenetic regulation, such as SR45, EIF4G, HD1 and ERF1-3 (Supplementary Fig. S6). Based on this finding, phosphorylation may modulate signals for the cell division of ECs and the differentiation of somatic embryos during sugarcane maturation by regulating proteins involved in epigenetic modifications.

The protein argonaute 1 (AGO1) was found to be regulated in EC14 and phosphorylated at T1049 and S1051 (which correspond to S1003 and S1005 in *Arabidopsis*, respectively). These phosphosites were identified in response to ABA treatment, which highlights the importance of the phosphorylation of these residues in stress responses (Wang et al., 2013). AGO1 also interacts with protein phosphatase 2C 32 (POL) during somatic embryo development (Fig. 4). POL was found to be unique in EC14 and phosphorylated at S192 (S189 in *Arabidopsis*), S208 (S205 in *Arabidopsis*) and S399 (not found in *Arabidopsis*). In *Arabidopsis*, POL is needed for maintenance of the dedifferentiated state of apical and root meristem cells during postembryonic growth by repressing CLAVATA (Song et al., 2020). Protein phosphatase 2C is an ABA coreceptor that negatively regulates ABA signaling by inhibiting SnRK2 (Saini et al., 2020). The phosphorylation of POL during the expression of embryogenic competence in ECs may be related to dephosphorylation of proteins that repress the differentiation of somatic embryos during maturation.

Chromatin reorganization is modulated by epigenetic mechanisms during somatic embryogenesis, including DNA methylation and posttranslational modifications of histone proteins (De-la-Pena et al., 2015; Kumar and Van Staden, 2017). The interactions among histone deacetylases and corepressors, such as TPL and TPR1, provides a link between the recognition of specific DNA sequences and chromatin remodeling activity (Plant et al.). In *Arabidopsis*, HD19 and TPL are recruited by APETALA2 (AP2) to repress the expression of target genes (Krogan et al., 2012). Here, putative histone deacetylase 19 (HD19, also known as histone deacetylase 1) was found to be up-regulated and phosphorylated at S420. This same phosphosite was previously identified in rice callus (Wang et al., 2017), and its corresponding site (S416) was found in suspensions of *Arabidopsis* cells (Nakagami et al., 2010). Both TPL and HD19 were also found to be phosphorylated during the induction of embryogenic competence in sugarcane and were associated with the control of gene expression during the acquisition of embryogenic competence (Fig. 4) (Almeida et al., 2020). However, unlike TPL and TPR1, HD19 was found to be up-regulated during the differentiation of sugarcane somatic embryos (Table 1). Thus, it can be hypothesized that HD1 plays a role in both the induction and manifestation of embryogenic competence.

Interactions among HOG1, SAM1, and SAM2 between maturation-related DAPs and DRPs was detected in this study (Fig. 4). In *Arabidopsis*, HOG1 is needed for DNA methylation-dependent gene silencing, an epigenetic mechanism of transcriptional repression (Rocha et al., 2005). HOG1 encodes an S-adenosylhomocysteine hydrolase (SAHH) responsible for producing methyl groups through the hydrolysis of S-adenosylhomocysteine, which is synthesized by S-adenosylmethionine synthetase (SAM) (Ouyang et al., 2012). These interactions in EC14 may be involved in methyl homeostasis and DNA methylation during the differentiation of sugarcane somatic embryos under maturation treatment. Further studies on polyamines can improve the available knowledge.

### 4.4. Other phosphoproteins with putative roles in somatic embryo differentiation in sugarcane

Ubiquitins are one of the marker proteins for different stages of somatic embryogenesis (Aguilar-Hernandez and Loyola-Vargas, 2018). In the present work, the interactions among proteins involved in protein modification, including UBC5, UBC35, SUMO1, and HD1, was observed in ECs under maturation (EC14) (Fig. 4). A phosphoproteomic study conducted with oil palm (*Elaeis guineensis*) suggested an E3 ubiquitin-protein ligase as a marker of somatic embryo development (Aroonluk et al., 2020). This finding indicates that protein ubiquitination may play a role in the differentiation and development of somatic embryos and suggests that ubiquitination can serve as a marker for the differentiation of sugarcane somatic embryos.

Interactions among and the regulation of CPK 4, 5, 6 and 20, APX3, HA1, and RHD2 were observed in EC14 cells (Supplementary Fig. S6). CPKs participate in signaling cascades involving Ca^2+^ as a secondary messenger and are important for development, growth, and stress responses (Shi et al., 2018). RHD2 is known to participate in the transient production of reactive oxygen species during *Arabidopsis* root hair development, and in these processes, Ca^2+^-dependent signaling and phosphorylation are important for its NADPH oxidase activity (Zhang et al., 2018). In contrast, APX3 contributes to homeostasis by eliminating ROS and using ascorbic acid as an electron donor (Narendra et al., 2006). HA1 is a proton pump located in the plasma membrane that is related to the embryogenic competence of sugarcane callus (Passamani et al., 2018). Thus, these data suggest that CPKs may be components of signal transduction pathways during the differentiation of sugarcane somatic embryos. Furthermore, signaling through ROS and Ca^2+^ appears to be fundamental for this process.

## 5. Conclusions

The integration of proteomics and phosphoproteomics data obtained during the differentiation of somatic embryos allowed us to observe the enhancement of and interactions among proteins and phosphoproteins involved in energy metabolism. Thus, it was possible to associate the regulation of proteins and phosphoproteins such as VHA-A/A1, RABA1c and RABA3, HOG1, SAM1 and 2, HD1, SUMO1 and UBC5, which are involved in the differentiation of somatic embryos. During the development of somatic embryos, activation of the photosynthetic apparatus was evidenced by the regulation of PSI-P, LHCB2.1, LHCB4.1 and LHCB6, and LHB1B2 and PSAE-1. In contrast, the maintenance of embryogenic competence in ECs during multiplication (EC0) is related to transcriptional control exerted by phosphoproteins with repressive activity, which inhibits the differentiation program. Thus, the maintenance of embryogenic competence includes the proteins and phosphoproteins TPL, ABF2, KIN10, LUH, SEU, SNRK2, NERD and GTB1. The identification of the unique phosphorylation motif [D-pS/T-x-D] during somatic embryo differentiation highlights protein phosphorylation as a central regulatory process during this process. The motifs [pS/T-D-x-E], [pS/T-P-x-R], [pS/T-F] and [pS/T-P] may also be involved in the maintenance and expression of embryogenic competence. These results reveal novel networks of protein regulation during somatic embryo differentiation and identify new players in *in vitro* morphogenesis.

## Acknowledgments

This research was supported by funds from the Fundação de Amparo à Pesquisa do Estado do Rio de Janeiro (FAPERJ) (Proc E26/211.690/2015; Proc E26/203.311/2017) and Conselho Nacional de Desenvolvimento Científico e Tecnológico (CNPq) (307755/2019-3), which were awarded to VS. This study was also financed in part by the Coordenação de Aperfeiçoamento de Pessoal de Nível Superior, Brasil (CAPES), Finance Code 001. Scholarships were provided by FAPERJ to LRX, FAA and VBP.

## CRediT authorship contribution statement

LRX, CSC and VS designed the research; FAA, LRX, LZP and VBP conducted the experiments; LRX, BPM, GASF, JJT and VS performed the proteomics and phosphoproteomics analyses; and all the authors read, reviewed and approved the manuscript.

