## Supplementary Figures for "Integrative proteomics and phosphoproteomics reveals phosphorylation networks involved in the maintenance and expression of embryogenic competence in sugarcane callus"

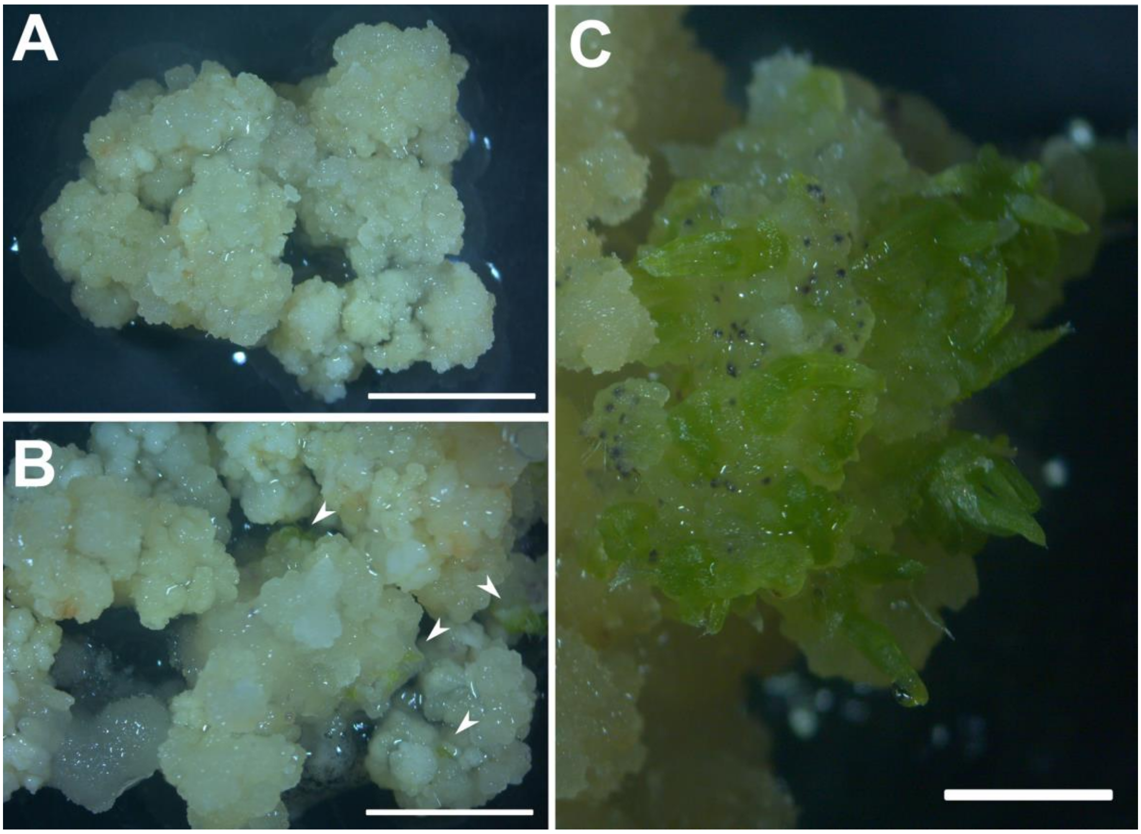


**Supplementary Fig. S1.** Morphological aspects of sugarcane embryogenic callus (EC) during the maturation of somatic embryos. ECs at the beginning of maturation treatment (EC0) (A). ECs after 14 days of maturation treatment (EC14) (B). Sugarcane somatic embryos formed after 42 days of maturation treatment (C). The arrowheads in (B) indicate the start of somatic embryo differentiation under maturation treatment. Bars in (A) and (B) = 5 mm. Bars in (C) = 2.5 mm.

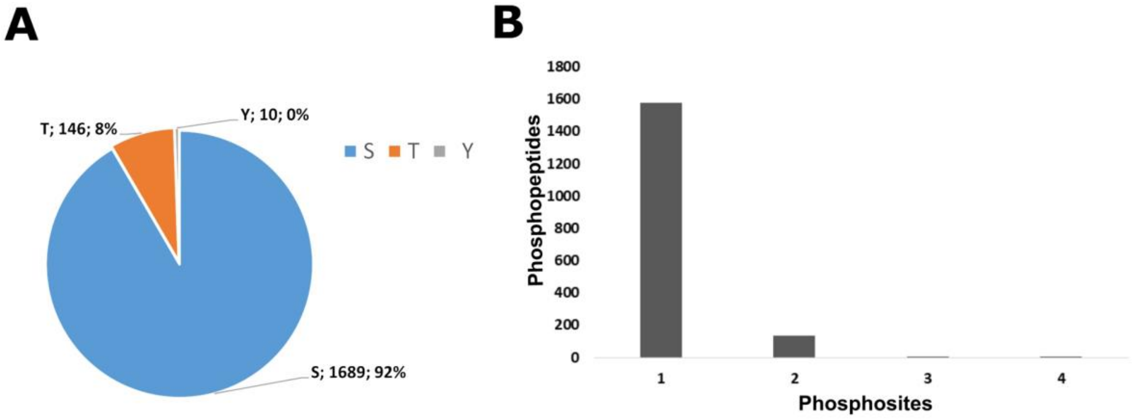


**Supplementary Fig. S2.** Features of the identified phosphopeptides. Proportion of phosphorylated serine, threonine and tyrosine amino acid residues (A). Number of phosphopeptides with 1, 2, 3 or 4 phosphosites (B).

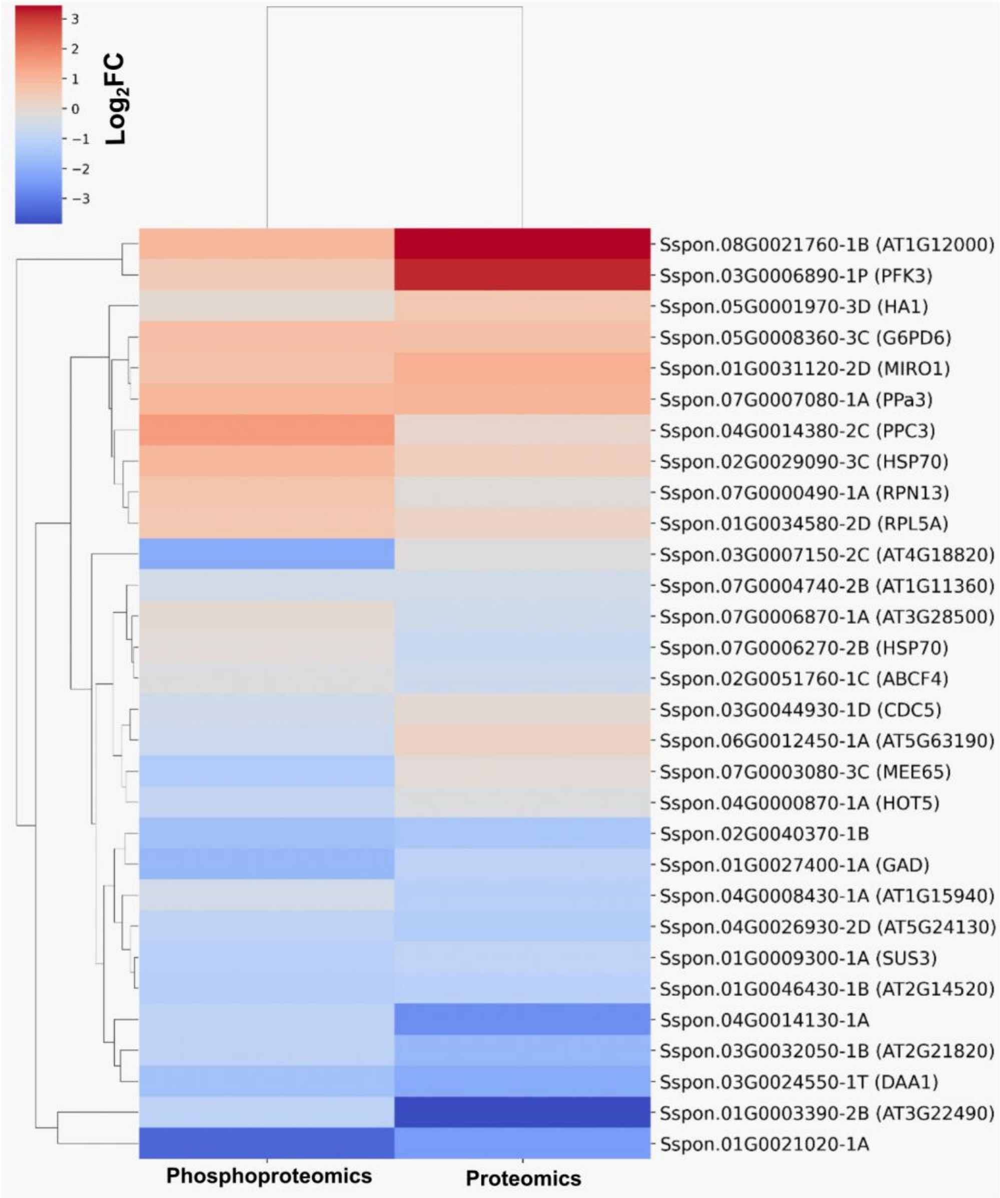


**Supplementary Fig. S3.** Heatmap showing proteins and phosphoproteins found in DAPs and DRPs identified in EC0 and EC14. The IDs were ranked according to the $Log_2$ FC values, and the *Arabidopsis thaliana* orthologs of the sugarcane proteins are shown in parentheses.

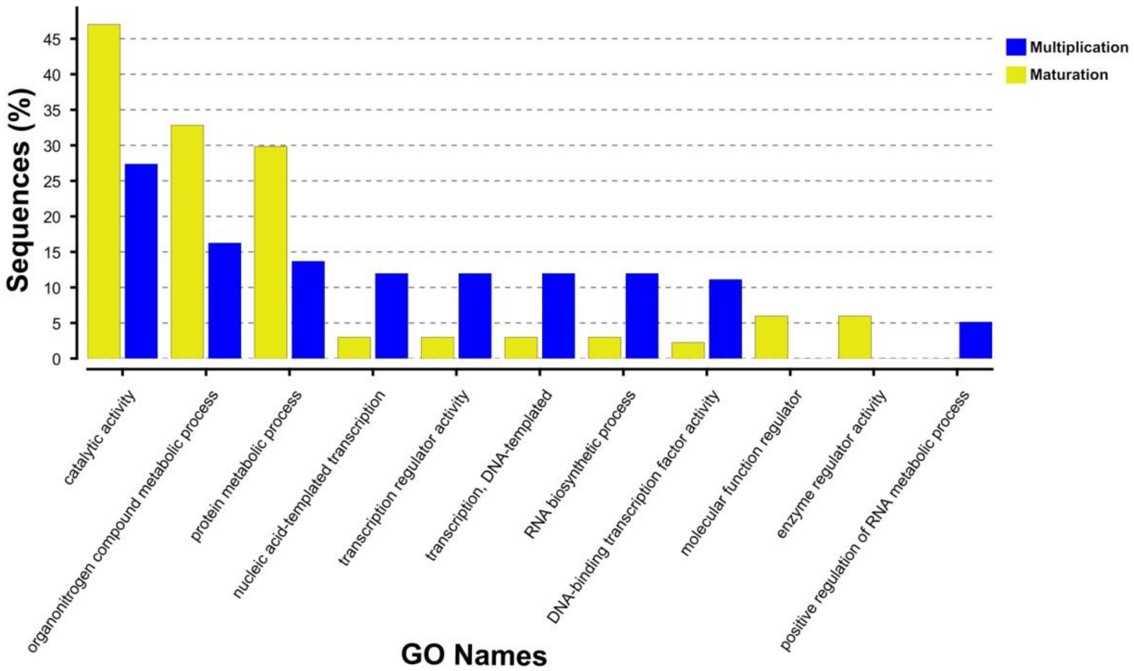


**Supplementary Fig. S4.** Gene ontology terms of biological processes, molecular functions, and cellular components enriched ($p < 0.01$) among the DRPs identified in the EC14/EC0 comparison. Only phosphoproteins with at least one phosphopeptide with a phosphorylation probability greater than 0.75 were included in the analysis.

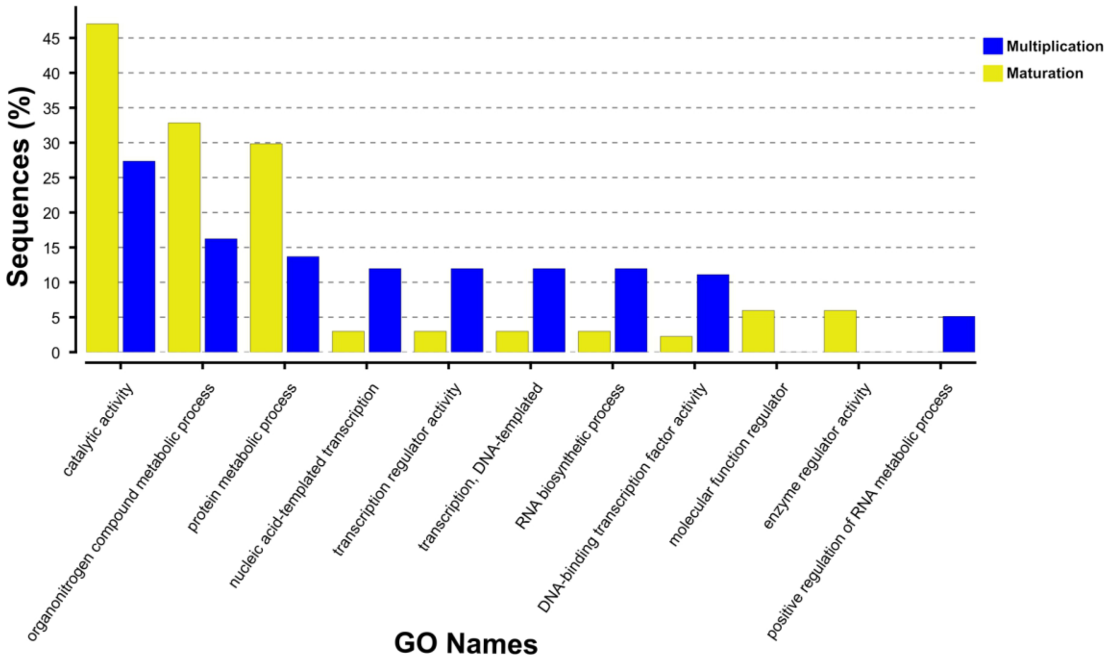


**Supplementary Fig. S5.** Gene ontology terms of biological processes, molecular functions and cellular components enriched among the DAPs identified in the EC14/EC0 comparison. OmicsBox software was used for gene ontology term enrichment analysis using Fisher's exact test ($p < 0.01$).

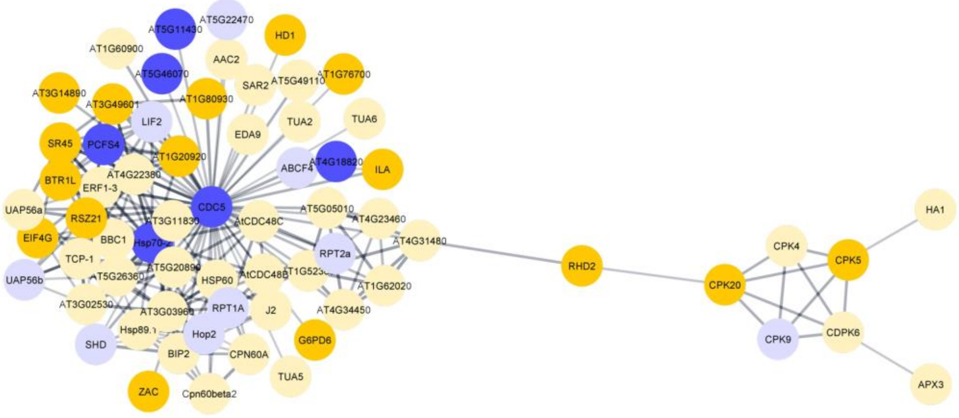


**Supplementary Fig. S6.** Protein-protein interaction networks between cyclin-dependent kinases and calcium-dependent kinases found among the DAPs and DRPs. The network was constructed using the *Arabidopsis* database in STRING after a search using the FASTA sequence of the sugarcane proteins.
